# 3D Organic Bioelectronics for Monitoring *In Vitro* Stem Cell Cultures

**DOI:** 10.1101/2022.03.30.486455

**Authors:** Achilleas Savva, Janire Saez, Chiara Barberio, Zixuan Lu, Chrysanthi-Maria Moysidou, Konstantinos Kallitsis, Aimee Withers, Charalampos Pitsalidis, Róisín M. Owens

**Affiliations:** Department of Chemical Engineering and Biotechnology, University of Cambridge, CB30AS Cambridge, United Kingdom; Microfluidics Cluster UPV/EHU, BIOMICs Microfluidics Group, Lascaray Research Center, University of the Basque Country UPV/EHU, Avenida Miguel de Unamuno, 3, 01006, Vitoria-Gasteiz, Spain; Ikerbasque, Basque Foundation for Science, E-48011 Bilbao, Spain; Department of Physics, Khalifa University of Science and Technology, P. O. Box 127788 Abu Dhabi, United Arab Emirates; Healthcare Engineering Innovation Center (HEIC), Khalifa University of Science and Technology, Abu Dhabi, United Arab Emirates

## Abstract

Three-dimensional *in vitro* stem cell models has enabled a fundamental understanding of cues that direct stem cell fate and be used to develop novel stem cell treatments. While sophisticated 3D tissues can be generated, technology that can accurately monitor these complex models in a high-throughput and non-invasive manner is not well adapted. Here we show the development of 3D bioelectronic devices based on the electroactive polymer poly(3,4-ethylenedioxythiophene)-poly(styrenesulfonate) - PEDOT:PSS and their use for non-invasive, electrical monitoring of stem cell growth. We show that the electrical, mechanical and wetting properties as well as the pore size/architecture of 3D PEDOT:PSS scaffolds can be fine-tuned simply by changing the processing crosslinker additive. We present a comprehensive characterization of both 2D PEDOT:PSS thin films of controlled thicknesses, and 3D porous PEDOT:PSS structures made by the freeze-drying technique. By slicing the bulky scaffolds we show that homogeneous, porous 250 um thick PEDOT:PSS slices are produced, generating biocompatible 3D constructs able to support stem cell cultures. These multifunctional membranes are attached on Indium-Tin oxide substrates (ITO) with the help of an adhesion layer that is used to minimize the interface charge resistance. The optimum electrical contact result in 3D devices with a characteristic and reproducible, frequency dependent impedance response. This response changes drastically when human adipose derived stem cells grow within the porous PEDOT:PSS network as revealed by fluorescence microscopy. The increase of these stem cell population within the PEDOT:PSS porous network impedes the charge flow at the interface between PEDOT:PSS and ITO, enabling the interface resistance to be extracted by equivalent circuit modelling, used here as a figure of merit to monitor the proliferation of stem cells. The strategy of controlling important properties of 3D PEDOT:PSS structures simply by altering processing parameters can be applied for development of a number of stem cell *in vitro* models. We believe the results presented here will advance 3D bioelectronic technology for both fundamental understanding of *in vitro* stem cell cultures as well as the development of personalized therapies.

## Introduction

Stem cells play a key role in regenerative medicine and tissue engineering due to their ability to proliferate, i.e. dividing and renewing themselves for long periods, and differentiate, i.e. commit to specific cell lineages.^1,2^ These abilities are sensitive to numerous chemical and physical cues induced by their environment.^3,4^ The development of *in vitro* systems that mimic the *in vivo* milieu has enabled a fundamental understanding of the effect of different cues^5^ and allowed development of novel stem cell based treatments.^6^ Stem cell research has recently benefitted enormously from advances in three-dimensional (3D) materials that are able to recapitulate tissue-like environments.^7^ Material properties such as surface texture, wettability, and macro-porous architecture impact the quality of these *in vitro* models.^8^

Three-dimensional (3D) organic bioelectronics — devices based on biocompatible, electrically active polymers — are proposed as versatile platforms to bridge the dimensionality mismatch between 2D/static electronics and 3D/dynamic biology.^9^ Electroactive scaffolds made from poly(3,4-ethylenedioxythiophene):poly(styrenesulfonate) (PEDOT:PSS) can be integrated into electrode or transistor configurations and allow electrical monitoring of 3D cell functions via conventional electrical measurements.^10^ These devices have been used to provide real-time information of the cell adhesion, growth and tissue formation^11^ as well as cellular protein conformation.^12^ Organic bioelectronic platforms with tissue-level complexity have also been demonstrated, e.g. a 3D model of the human intestine,^10^ and show potential as accurate animal alternatives for disease modelling, drug discovery and tissue engineering.^13^

PEDOT:PSS water dispersions allow facile solution processing to form highly biocompatible 3D porous structures with metal-like conductivity and mechanical properties approaching human tissue. The freeze-drying method (also known as ice-templating or lyophilization)^14^ is widely used to form 3D structures with controlled porous size and architecture.^15^ Processing parameters such as the solution concentration, freezing rate, or the use of additives^16,17^ can be used to fine-tune the morphology and the microstructure of the scaffold. Among these parameters, the use of crosslinking agents is essential for water-stability of PEDOT:PSS-based 3D structures. 3-glycidyloxypropyl)trimethoxysilane (GOPS) is the most commonly used crosslinker^18^ and leads into PEDOT:PSS structures with adequate stability in cell-culture conditions for a variety of bioelectronic applications.^19,20^ Alternative crosslinking strategies involve the use of poly(ethylene glycol)diglycidyl ether (PEGDE)^21^ and divinyl-sulfone (DVS).^22^ Both GOPS and PEGDE render PEDOT:PSS water-stable via similar mechanisms — a reaction of the epoxy rings moiety present in their structures with the weakly nucleophilic PSS^-^.^23^ Importantly, the choice of cross-linker impacts the electrochemical,^24^ surface topography,^21^ and mechanical properties^25^ of PEDOT:PSS structures. Considering these properties, PEDOT:PSS-based scaffolds have come to the fore as multifunctional biomaterials with tailor-made properties allowing the development of smart bioelectronic interfaces for *in vitro* models.^23^

Organic bioelectronic interfaces for in vitro 3*D* stem cell cultures e can be customized to mimic accurately the micro-environment required for stem cell growth. For example, Iandolo et al. developed composite PEDOT:PSS/collagen scaffolds with highly elastic mechanical properties that supported “soft” neural crest-derived stem cell culture.^26^ In addition, highly porous PEDOT:PSS scaffolds with more rigid mechanical properties have been proposed for the development of bone tissue.^27^ However, there is a need to develop noninvasive analysis techniques to monitor the behavior of stem cell tissue in real time during *in vitro* culturing. Although, recent advancements show the development of non-invasive technique to monitor stem cell growth,^28,29^ technology that can accurately assess the functionality of these complex models in a high-throughput and dynamic manner would further advance stem cell research towards commercialization.

Here we show the development of 3D bioelectronic devices based on PEDOT:PSS that are used for non-invasive, electrical monitoring of stem cell culture. We propose the use of electroactive 3D PEDOT:PSS structures as hosts of human adipose derived stem cells (hADSCs). We show that the electrical, mechanical and wetting properties as well as the pore size/architecture of these 3D PEDOT:PSS structures, produced with the freeze drying technique, can be fine-tuned simply by changing the processing crosslinker. PEDOT:PSS structures crosslinked with PEGDE exhibit increased electrical conductivity, volumetric capacitance, and water retention as well as being more elastic compared with PEDOT:PSS structures crosslinked with GOPS. The pore size and architecture are strikingly different for scaffolds made PEDGE and GOPS as well as well mixtures of PEGDE and GOPS, as revealed by scanning electron microscopy (SEM). PEDOT:PSS scaffold slices (250 μm thick) were attached to an Indium-Tin oxide substrate (ITO) with the help of an adhesion layer – a PEDOT:PSS:GOPS thin film that is used to molecularly attach the scaffold slice, minimizing the interface charge resistance, and enabling a characteristic frequency dependent impedance response. hADSCs were seeded directly in the scaffold slice and shown to adhere, survive and fully colonize the scaffold after 10 days as revealed by live/dead assay and fluorescence microscopy. Electrochemical impedance spectroscopy measurements are used to monitor cell proliferation - while the cells are growing, a progressive increase in the impedance magnitude is observed. This physical process is simulated with an interface resistance in the proposed equivalent circuit model and used as a figure of merit to monitor cell proliferation

## Results and discussion

We first studied the electrochemical and surface properties of PEDOT:PSS thin films crosslinked with GOPS (3 wt%) and PEGDE (3 wt%). Both GOPS and PEGDE render PEDOT:PSS insoluble in electrolytes, via a reaction between the ring epoxide moiety, present in both crosllinker’s structure, with the PSS^-^ chains (Figure 1a). However, we found that PEDOT:PSS films crosslinked with PEGDE are more hydrophilic compared with PEDOT:PSS crosslinked with GOPS. As shown in figure 1b, the water contact angle was measured at 62 ° on PEDOT:PSS cross-linked with GOPS and at 7 ° on PEDOT:PSS cross-linked with PEGDE. Although GOPS crosslinked PEDOT:PSS films have been shown to be excellent interfaces for cell culture, the enhanced hydrophilicity of PEGDE crosslinked PEDOT:PSS can further facilitate cell attachment and growth, in line with previously reported studies.^21 30 31^

**Figure 1:**
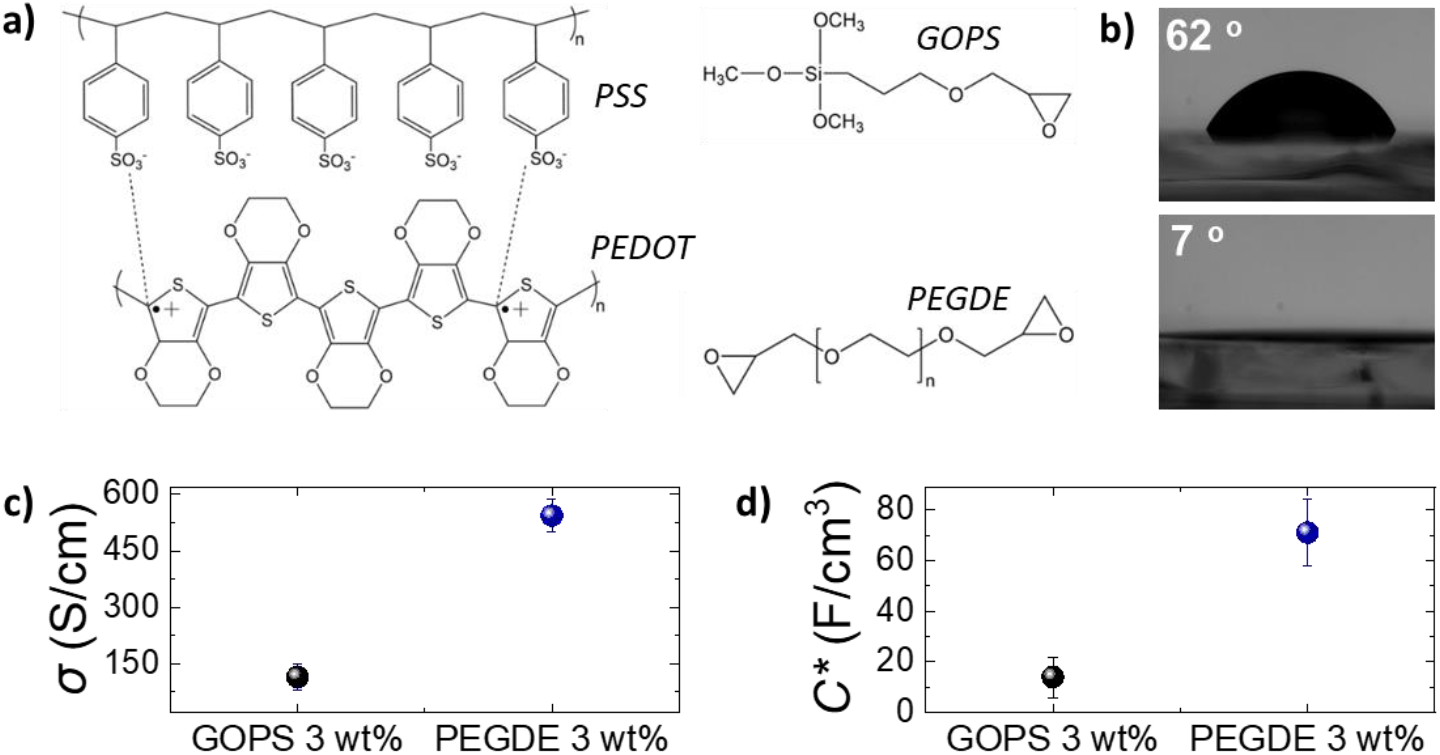
**a)** The chemical structures of the conducting polymer PEDOT:PSS and the crosslinkers (3-glycidyloxypropyl)trimethoxysilane (GOPS) and poly(ethylene glycol)diglycidyl ether. **b)** Digital pictures of the water contact angle on top of a PEDOT:PSS film crosslinked with GOPS 3 wt% (top) and with PEGDE 3 wt% (bottom). **c)** Electrical conductivity (*σ*) and **d)** volumetric capacitance (C*) of PEDOT:PSS films crosslinked with GOPS 3wt% and PEGDE 3 wt% (N=6).

To accurately characterize the electrochemical properties of PEDOT:PSS films, we used micro-fabricated organic electrochemical transistors (OECTs) and electrodes. By using current-voltage measurements of PEDOT:PSS OECT channels of controlled thickness (Figure S1), we calculated the electrical conductivity of PEDOT:PSS films crosslinked with PEGDE (3 wt%) at 544 S/cm and with GOPS (3 wt%) at 125 S/cm (Figure 1c). Furthermore, by using electrochemical impedance spectroscopy (EIS) measurements (Figure S1), we found that the volumetric capacitance of PEDOT:PSS films crosslinked with PEGDE (3 wt%) is 71 F/cm^3^ and with GOPS (3 wt%) is 14 F/cm^3^. These results show that PEGDE crosslinked PEDOT:PSS thin films exhibit superior electrochemical properties compared with GOPS crosslinked PEDOT:PSS thin films. However, we should note that the typical PEDOT:PSS crosslinking formulation that is used for thin bioelectronic devices is GOPS (1 wt%),^20^ with electrical conductivity values in the range of 400 S/cm^24^ and volumetric capacitance of 39 F/cm^3^.^32^ We found that the OECT transconductance (*g*_m_) is higher for PEDOT:PSS channels crosslinked with PEGDE (3 wt%) compared with PEDOT:PSS channels crosslinked with the standard GOPS (1 wt %) (Figure S2). Overall, PEDOT:PSS crosslinked with PEGDE (3 wt%) shows superior electrochemical properties compared with both GOPS (3 wt%) and GOPS (1 wt%) crosslinked PEDOT:PSS films. However, it is important to mention that PEGDE crosslinked PEDOT:PSS thin films delaminate from the glass substrates after a few days of operation in cell culture conditions (Figure S3). We attribute this to the absence of silanes groups in the PEGDE molecule (as opposed to GOPS), which react with glass substrates and render PEDOT:PSS strongly attached on a glass support for long-term.^18^ Although we believe the PEGDE crosslinking strategy can be potentially used for the development of more sensitive thin film bioelectronic devices, further optimization of thin film devices is beyond the scope of this study.

We then fabricated PEDOT:PSS-based porous scaffolds crosslinked with different concentrations of GOPS and PEGDE *via* the freeze-drying technique.^11,27^ Both GOPS and PEGDE result in robust and stable 3D structures that maintained their shape for more than six months immersed in aqueous electrolytes (Figure S4). However, we found that the choice of the crosslinker regulates scaffold properties that are important for cell growth such as, water retention (Figure 2a), elasticity (Figure 2b) and pore size (Figure 2c). The water retention ability of 3D scaffolds used as hosts for tissue growth is an essential property that regulates cell media penetration and impacts cell proliferation. As shown in figure 2a, we found that PEDOT:PSS scaffolds crosslinked with GOPS (3 wt%), PEGDE (1 wt%) and PEGDE (3 wt%) retain substantial amounts of water - mean values 1712 %, 1744 %, 1777 %, respectively. In contrast, the water retention ability is significantly reduced to a mean value of 327 % and 213 % for scaffolds crosslinked with PEGDE (5 wt%) and PEGDE (10 wt%), respectively (Figure S5). We also measured the mechanical properties of PEDOT:PSS scaffolds crosslinked with both GOPS (3 wt%) and PEGDE (3 wt%) and calculated a mean Young’s modulus of 32.2 KPa and 17.9 KPa, respectively (Figure 2b). These results show PEDOT:PSS scaffolds crosslinked with PEGDE are more elastic compared with those crosslinked with GOPS.^23^ The molecular structures and mechanisms of crosslinking of both GOPS and PEGDE can explain these observations: In GOPS crosslinked PEDOT:PSS, PSS^-^ chains are interconnected with an intermediate/rigid silyl ether bond formed between two GOPS molecules.^18^ In contrast, PEGDE crosslinked PEDOT:PSS, PSS^-^ chains are directly interconnected with a single PEGDE molecule,^21^ which is expected to lead in more flexible bonds and as a results, lower Young’s modulus.

**Figure 2:**
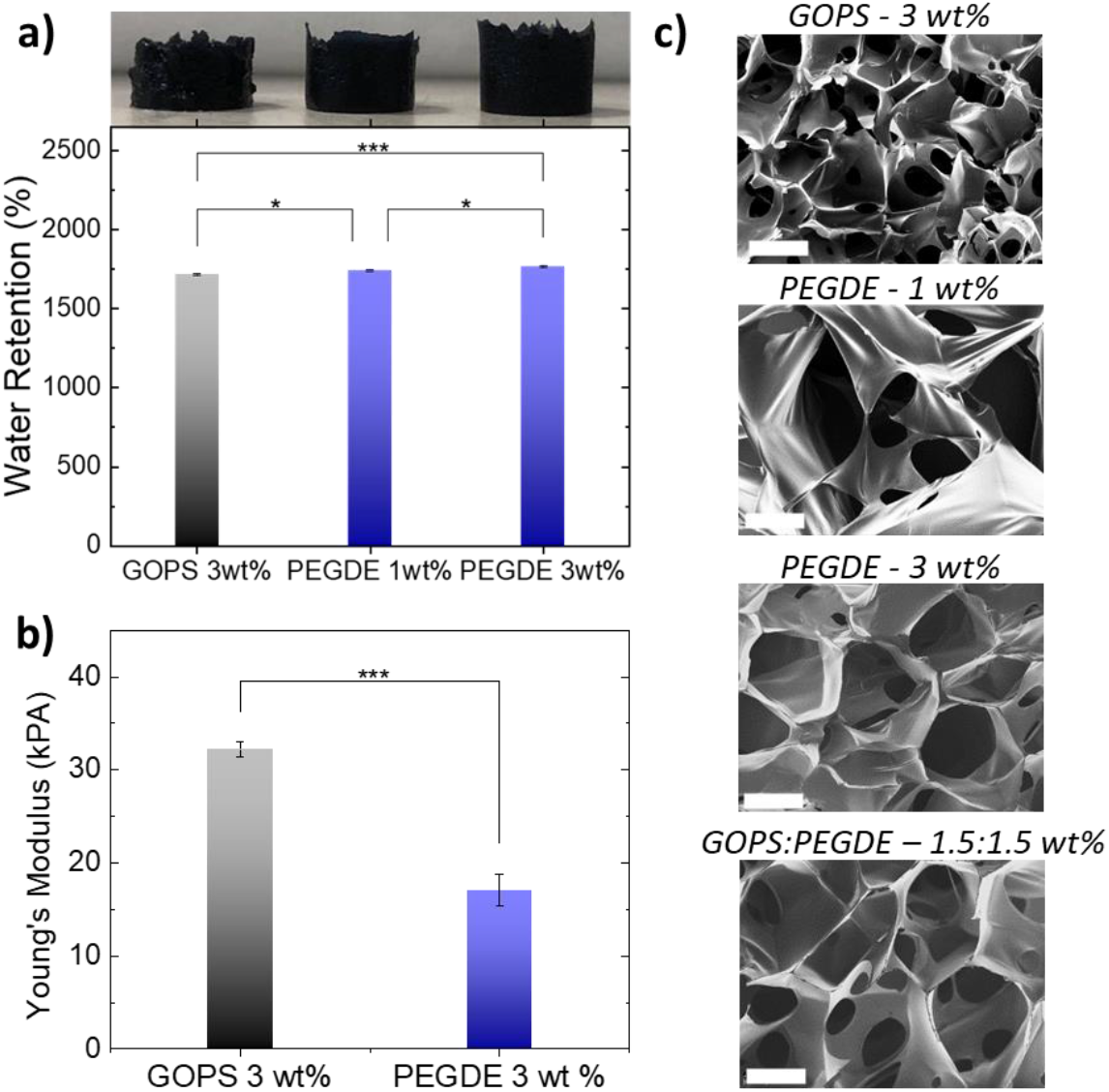
**a)** Water retention ability of PEDOT:PSS-based scaffolds crosslinked with GOPS 3 wt%, PEGDE 1 wt% and PEGDE 3 wt%. The digital pictures on top of the graph show the scaffolds for the corresponding concentration of crosslinker after been swollen in DI water (N=6). **b)** Young’s moduli for PEDOT:PSS scaffolds crosslinked with GOPS 3 wt% and PEGDE 3 wt% extracted from compression tests (N=6). **c)** Scanning electron microscopy measurements of scaffolds prepared with different crosslinkers. Scale bars = 40 μm.

Moreover, as shown in figure 2c, we examined the pore morphology of all the different PEDOT:PSS-based scaffolds using scanning electron microscopy (SEM). PEDOT:PSS scaffolds crosslinked with GOPS (3 wt%) exhibit an average pore size in the range of ∼50 μm, with anisotropic pore architecture, in agreement with previously reported studies. In contrast, PEDOT:PSS scaffolds produced with PEGDE show larger average pore size, in the range of 100-200 μm. Scaffolds made with PEGDE (3 wt%) show more structured, honeycomb-like architecture compared with PEGDE (1 wt %). When the concentration of PEGDE crosslinker is further increased to more than (5 wt%), the scaffolds show no porosity (Figure S5), which can also explain the significant reduction of water retention observed in figure 2a for such high PEGDE concentrations. Importantly, we also used a mixture GOPS and PEGDE crosslinkers to produce PEDOT:PSS 3D structures, with interconnected porous network of different sizes. The above-mentioned observations are also visible in larger area SEM images shown in figure S6. Overall, all the PEDOT:PSS-based scaffolds produced with the different crosslinkers, concentrations and mixtures (except PEGDE higher than 5 wt%) exhibit highly open anisotropic pore architecture with average pore sizes suitable for cell penetration.

We then integrated these PEDOT:PSS-based porous structures into 3D bioelectronic devices. We chose to proceed with PEDOT:PSS crosslinked with PEGDE (3 wt%) for two main reasons. First, the superior electrical properties of PEGDE crosslinked PEDOT:PSS (Figure 1) could lead to devices with higher sensitivity. Second, the mechanical properties, combined with larger pore size of PEGDE crosslinked PEDOT:PSS scaffolds are more suitable for human adipose derived stem cell (hADSC) cultures. As shown in Figure S7, individual stem cells grown in standard well plates show elongated shape with sizes ranging between 200 μm and 400 μm, in line with previously published studies.^33^

The fabrication process of the 3D bioelectronic devices we used is shown in Figure 3a and 3b. First, the PEDOT:PSS scaffolds were sliced using a vibratome to produce scaffolds slices of controlled thickness (i.e. 250 μm) and homogeneous pore size distribution (Figure 3a). These highly porous slices were then attached on Indium-Tin Oxide (ITO) transparent conducting substrates with the help of an adhesion layer — a PEDOT:PSS thin film with GOPS crosslinker used to molecularly “hook” the PEDOT:PSS scaffold slice. The device was finalized with the attachment of a circular mask (i.e. Kapton® tape - diameter = 7 mm) used to define the device active area at 0.39 cm^2^ followed by the attachment of a glass well to contain cells and cell media as it will be discussed later.

**Figure 3:**
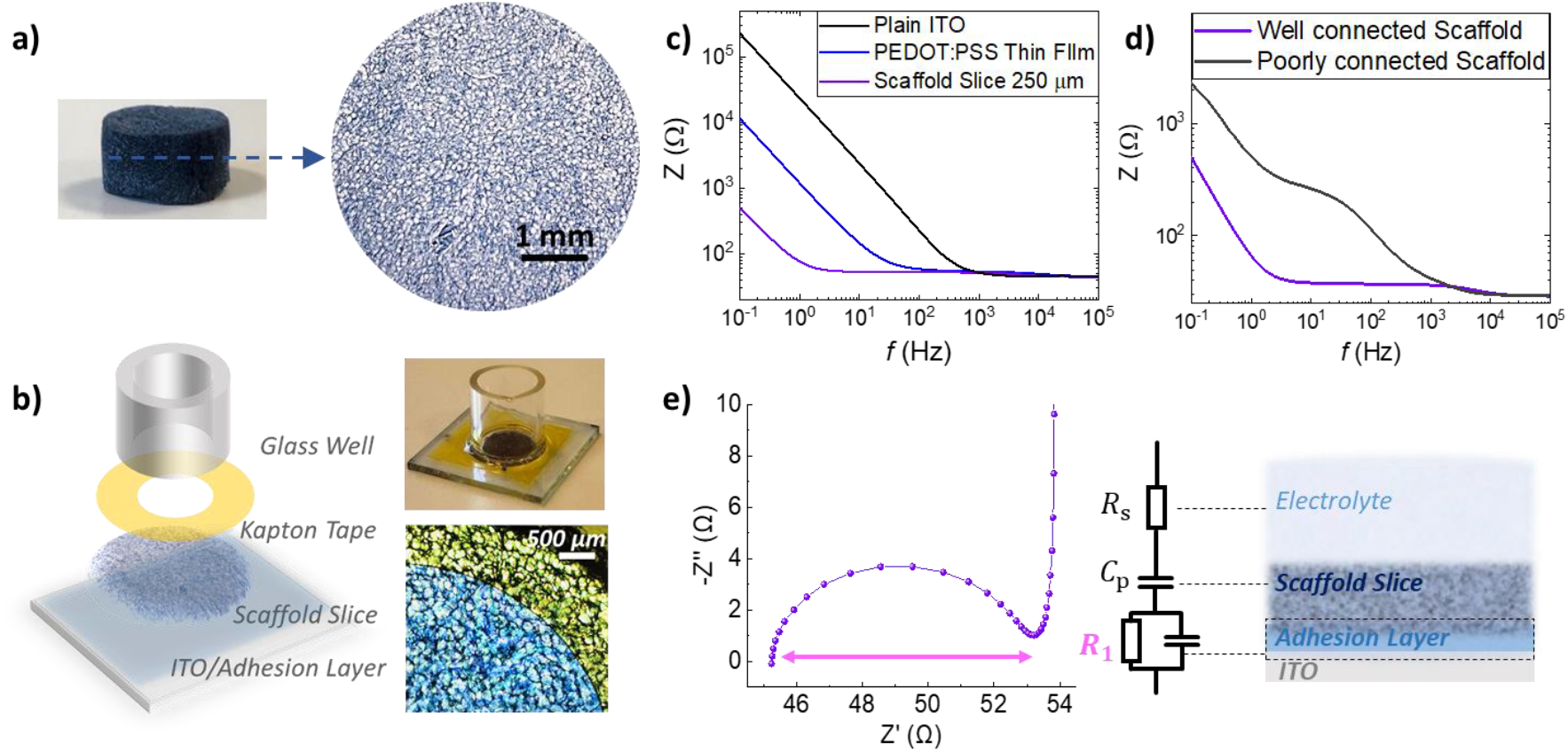
**a)** Digital pictures of a PEDOT:PSS scaffold and a 250 μm thick scaffold slice made of PEDOT:PSS crosslinked with PEGDE 3 wt%. **b)** Schematic of the 3D bioelectronic device developed in this study and as shown in the digital picture on the top right. A top view of the edge of this device is shown in the bottom left microscope image. The electrochemical area of the device is well defined by the micro-patterned Kapton tape. **c)** Impedance magnitude versus frequency plots for the devices bearing the same electrochemical area as defined by the Kapton tape with plain ITO (black), ITO/adhesion layer (blue) and with the scaffold slice attached (purple). **d)** A typical impedance versus frequency plot obtained for a well-connected scaffold slice (of ITO/adhesion layer/scaffold slice devices-purple) and for a poorly connected scaffold (no use of adhesion layer - grey). **e)** A typical Nyquist plot of an optimized ITO/adhesion layer/scaffold slice device and the equivalent circuit model used to raw EIS data.

As shown in figure 3b, the impedance magnitude is compared for plain ITO devices, ITO/adhesion layer devices and ITO/adhesion layer/scaffold slice devices of the same area (0.39 cm^2^). We observe a striking drop of the impedance magnitude at low frequencies (i.e. ∼0.1 Hz - 100 Hz) for the ITO/adhesion layer devices compared with the plain ITO devices and a further decrease of the impedance magnitude for the ITO/adhesion layer/scaffold slice devices. Since PEDOT:PSS capacitance scales with volume,^34–36^ the striking drop of the impedance magnitude in the low frequency regime can be attributed to the electrical connection of the 250 μm thick porous PEDOT:PSS scaffold slice with the conducting ITO substrate. To quantify these changes, we calculated the overall capacitance of the system at 0.1 Hz to be 168 μF/cm^2^, 358 μF/cm^2^, 7692 μF/cm^2^ for plain ITO, ITO/adhesion layer and ITO/adhesion layer/scaffold slice devices, respectively. Here we should note that despite the good approximation of the scaffold capacitance per unit area, a more accurate estimation should involve the scaffold thickness, average pore size and number of pores per unit area. Nevertheless, the above results prove that the scaffold is well electrically connected on the ITO substrate and promotes the importance of the adhesion layer. As shown in figure 3d, a scaffold slice that was simply mechanically pressed on the ITO substrate (named as a poorly connected scaffold) shows high impedance in frequencies below 1000 Hz. In contrast, when an adhesion layer is used the impedance magnitude is significantly smaller at this frequency range. The latter can be attributed to a good electrical connection of the scaffold slice on the ITO substrate. The characteristic semicircle of the 3D devices can be seen in the Nyquist plot graph in figure 3e, together with the equivalent circuit model used to fit the experimental data. We used a resistor element for the electrolyte resistance (*R*_S_), in series with a capacitor for PEDOT:PSS (*C*_P_), and with an *R*_1_-*C*_1_ circuit element for the crucial PEDOT:PSS/ITO interface. The quality of this interface is crucial for charge extraction from PEDOT:PSS to the ITO and therefore, *R*_1_ can be used as a figure of merit to describe the charge transfer at this interface. As shown in Figure 3e, *R*_1_ can be easily calculated by the absolute magnitude of the semicircle formed in the Nyquist plot representation of impedance data (i.e. subtraction of the absolute values that the semicircle intersects the x-axis). Based on all the above, the use of the adhesion layer between the ITO and the PEDOT:PSS scaffold slice, drastically improves charge collection and minimizes *R*_1_ between 5-20 Ohm as calculated from more than 10 individual devices.

The establishment of a reproducible and characteristic impedance spectra allowed the monitoring of cell-related changes with impedance measurements. The transparency of the ITO substrates together with the high PEDOT:PSS scaffold porosity also allowed the monitoring of cell growth via confocal microscopy. As shown in figure 4a-d, fluorescence images obtained from a viability assay proved that a healthy stem cell culture developed within the 3D PEDOT:PSS-based devices 10 days after seeding. We observed (Figure 4a,b) that stem cells proliferate and colonize the scaffold forming dense cultures at day 10 after seeding (figure 4b). Regimes where cells grow around the PEDOT:PSS scaffold pores can be also seen in figure 4c, and stem cell network that is developed in all direction and on different planes can be seen in figure 4d. From these images, we also observe that stem cells grown within the PEDOT:PSS scaffold preserve a similar elongated shape compared with the cells grown in control well-plates (figure S7). These results also prove that PEDOT:PSS-PEGDE scaffolds slices and all the other materials used for the development of the 3D devices are cyto-compatible.

**Figure 4:**
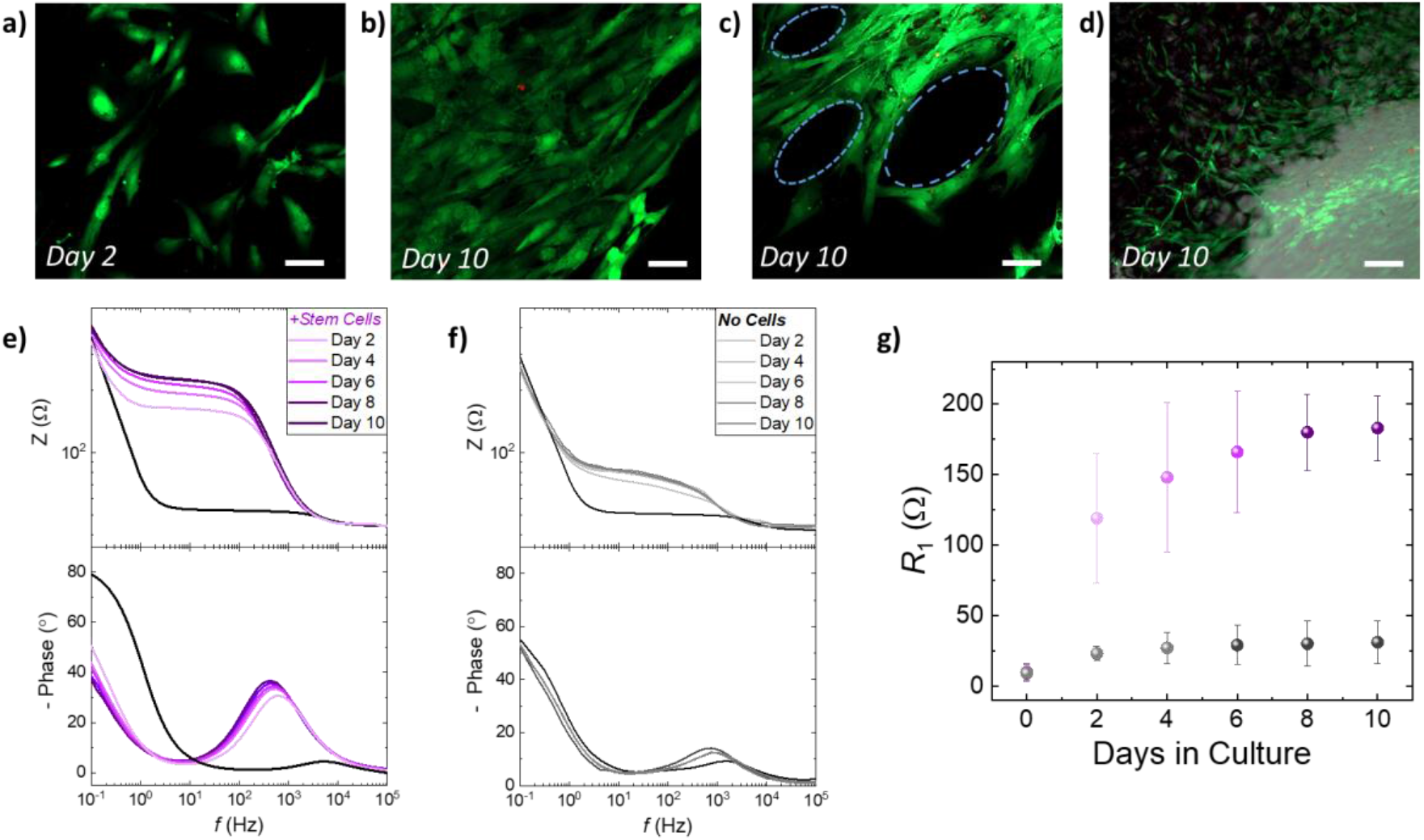
**a)-d)** Live/Dead assays of human adipose derived stem cells cultures grown within the proposed ITO/adhesion layer/scaffold slice bioelectronic devices, comprising a PEDOT:PSS scaffold crosslinked with PEGDE 3 wt%. The impedance magnitude and the phase of the impedance obtained every 2 days for overall 10 days for identically prepared devices: **e)** that have not been seeded with hADSCs and **f)** seeded with hADSCs. **g)** the interface resistance (*R*_1_) extracted as a figure of merit and plotted as a function of days monitoring the stem cell tissue growth for control devices (black circles) and for devices that have been seeded with hADSCs (purple) (N=5).

Importantly, the electroactive properties of the PEDOT:PSS-based scaffold slices were leveraged to monitor the increase in stem cell population using impedance measurements. As shown in figure 4e, we monitored the impedance spectra changes of devices that have been seeded with hADSCs for 10 days. The impedance magnitude and the phase of the impedance showed a distinct increase at the frequency range ∼ 1 Hz - 1000 Hz over the period of 10 days in culture (figure 4e). These changes are also distinct in the characteristic semicircle of the Nyquist plots (discussed earlier in figure 3) as shown in figure S8. In contrast, the impedance magnitude, and the phase of the impedance of identically prepared devices that have not been seeded with stem cells show only a slight increase in the range of frequency ∼ 1 Hz - 1000 Hz (figure 4f). We applied the equivalent circuit model and extracted *R*_1_ as the figure of merit of our system, as discussed earlier in figure 3e. As expected from the raw impedance data (figures 4e, 4f and S8), *R*_1_ drastically increases for the devices seeded with hADSCs (from 9 Ohm before seeding to 119 Ohm at day 2 to 183 Ohm at day 10). The devices that have not been seeded with hADSCs show slight increase in *R*_1_ during the 10 days period of incubation (from 10 ohm in day 0 to 31 Ohm at day 10). We suggest that the increase in *R*_1_ can be correlated with the increase in stem cell population within the PEDOT:PSS scaffolds. Our optical measurements show that stem cells proliferate and colonize scaffolds in all directions — an event that can disrupt the electrically conducting polymer network and increase the impedance of the system. Moreover, the inclusion of stem cells within the pores of PEDOT:PSS structure affects the crucial interface between the PEDOT:PSS scaffold slice and the ITO surface and impedes the charge transfer. Therefore, our proposed devices serve as a facile platform to monitor stem cells proliferation with conventional electrical measurements.

## Conclusion

In summary, we showed the development of PEDOT:PSS-based electroactive porous scaffolds that are used as hosts for human stem cell cultures. We demonstrated that the mechanical, electrical and morphological properties of these scaffolds can be tailored by simply changing the crosslinking parameters, a significant advantage over other scaffold materials that are commonly used. PEDOT:PSS crosslinked with PEGDE shows excellent electrical conductivity, volumetric capacitance as well as hydrophilic properties. 3D porous structures made by this formulation *via* lyophilization, showed increased water retention ability, more elastic behavior and larger pore size distribution compared with the established GOPS crosslinking strategy.

Importantly we demonstrated that the porous network can be controlled by using mixtures of GOPS:PEGDE crosslinkers – a feature that can be exploited to tailor scaffold properties for specific cell cultures. Furthermore, we reported the development of 3D bioelectronic devices with characteristic and reproducible electrochemical impedance signal. The use of an adhesion layer — a PEDOT:PSS thin film with GOPS crosslinker used to molecularly “hook” the PEDOT:PSS scaffold slice on ITO substrates — minimizes the interface resistance (*R*_1_) between the scaffold slice and the ITO substrate. These devices supported the adhesion and growth of viable hADSCs cultures over 10 days, as revealed by viability assays. The excellent electrical properties of the device and the PEDOT:PSS scaffolds are utilized to monitor stem cell growth with non-invasive impedance measurements. The device impedance signature signal changes upon stem cell proliferation which is correlated with *in situ* fluorescence imaging of the stem cells tissue. This change is quantified by calculating the increase of the interface resistance caused by inclusion of the cells in the scaffold’s pores. In conclusion, the strategy of controlling important properties of 3D PEDOT:PSS structures shown here, can be applied for development of a number of stem cell *in vitro* models. Further development of these platforms can also have added features, such the ability to stimulate stem cell cultures aiming in controlled cell-fate direction. We believe the results presented here pave the way for further advancements in 3D bioelectronic technology and specifically for 3D stem cell research for both fundamental understanding of in vitro stem cell cultures as well as the development of personalized therapies.

## Experimental

### Materials

PEDOT:PSS Clevios PH - 1000 was purchased from Heraeus GmbH. (3-glycidyloxypropyl)trimethoxysilane - GOPS (product no. 440167), poly(ethylene glycol) diglycidyl ether -PEGDE (product no. 475696), ethylene glycol (product no. 324558), sodium dodecylbenzenesulfonate - DBSA(product no. 44198) and glass wells (product no. CLS316610) were purchased from Sigma-Aldrich.

### Contact angle measurements

Water contact angles were measured on thin films of PEDOT:PSS with a KRUSS DSA 100E goniometer by using the sessile drop method. All measurements were taken 5 seconds after the water drop deposition on each films in a cleanroom environment with constant relative humidity of 45% following the recommendation by C. Duc et al.^37^

### 2D device micro-fabrication

The fabricated device were OECT arrays with the PEDOT:PSS. One row with three OECTs had 50 μm by 50 μm channels, and for the other row, the three OECTs had 100 μm by 100 μm channels. To fabricate these OECTs, 4 inch glass wafers first were cleaned by sonication in an acetone and then Isopropanol 15 minutes. The wafers were rinsed with DI water and baked 15 min at 150 °C. To pattern for contact tracks, a negative photoresist, AZ nLOF2035 (Microchemicals GmbH) was spun on the glass wafer with 3000 rpm for 45s and exposed with UV light using mask aligner (Karl Suss MA/BA6). The photoresist was developed in AZ 726 MIF developer (MIcroChemicals) developer for 28 s. Ti (5 nm)/Au (100 nm) layer as conductive track was deposited by e-beam evaporation on top of wafer and the Ti-Au metal layer was lifted-off by soaking in NI555 (Microchemicals GmbH) overnight. Prior to the deposition of 2 μm layer (sacrificial layer) of parylene C ((SCS), the wafer is soaked with 3% A174 (3-(trimethoxysilyl)propyl methacrylate) in ethanol solution (0.1% acetic acid in ethanol) for 60 seconds to promote the parylene C adhesion on the wafer. An anti-adhesive layer of Micro-90 in DI water (2% v/v solution) was spun (1000 rpm for 45 seconds), and then the second layer of 2 μm parylene C (SCS) was deposited. A layer of positive photoresist AZ 10XT (Microchemicals GmbH) was spun with 3000 rpm for 45s and developed in AZ 726 MIF developer (MIcroChemicals) for 6 min to pattern OECT channel. Reactive ion etching (Oxford 80 Plasmalab plus) opened the window for deposition of either Clevios PH1000 PEDOT:PSS (Heraeus). The PEDOT:PSS solution containing 5 vol% ethylene glycol, 0.26 vol% dodecylbenzenesulfonic acid (DBSA),and 1-3 vol% (3-glycidyloxypropyl)trimethyloxy-silane (GOPS) or 1-3% PEGDE was spin-coated at 3000 rpm for 45s. The samples were baked at 90 °C for 1 min, and the sacrificial layer was peeled off. Finally, the sample was put on a hot plate at 130 °C for 1 h before use.

### 2D device characterization

OECTs were characterized using a dual-channel source-meter unit (NI-PXI) with custom-written control code in LabVIEW. All measurements were performed using an Ag/AgCl pellet (D = 2 mm × H = 2 mm – Warner instruments) as the gate electrode. PBS 1X electrolyte was placed in a PDMS well on top of the OECTs at a constant volume (150 μL). The frequency dependent response of the transconductance was extracted using a method described previously by Rivnay et al.^2^ Electrochemical impedance spectroscopy (EIS) measurements were performed using a PG128N Metrohm-Autolab potentiostat, with a standard three-electrode setup in PBS_(aq.)_ 1 X. The films were addressed as the working electrode, while a Pt mesh and Ag/AgCl were used as the counter and reference electrode, respectively. EIS was performed at a frequency range between 100 kHz to 0.1 Hz at *V* = 0 V *vs V*_OC_ with an AC amplitude of 10 mV. We extracted the volumetric capacitance value for each system by determining the value of capacitance at 0.1 Hz using equation 1 and dividing it with the film volume (area x thickness):

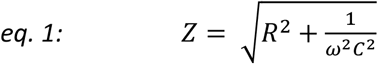

### PEDOT:PSS macro-porous scaffolds and slices

Scaffolds were fabricated from an aqueous dispersion of PEDOT:PSS (Clevios PH-1000, Heraeus) at a concentration of 1.2 wt%, using the freeze drying process.^11^ The different PEDOT:PSS solution formulations were prepared as follows: 5 ml of PEDOT:PSS PH1000 solution was first filtered with 0.8 μm cellulose acetate filters. Then 26 μl of DBSA (0.5 wt%) and different amount of crosslinkers, i.e. 150 μl of GOPS (3 wt%), PEGDE 50 – 500 μl (1-10 wt%) or a mixture of 75 μl GOPS and PEGDE 75 μl (1.5:1.5 wt%) were added in the solution followed by sonication at room temperature for 10 minutes. The different PEDOT:PSS solution formulations were poured into either 24, 48 or 96 well-plates that were used as molds for producing macro-porous scaffolds of different dimensions. The well-plates were placed in the freeze-dryer (Virtis AdVantage 2.0 BenchTop from SP Scientific) and frozen from 5 to −30 °C at a controlled cooling rate, after which point the ice phase was sublimed from the scaffolds. The samples were then removed and dried at 80 °C for 4 h and. PEDOT:PSS scaffold slices of controlled thickness were produced by slicing the bulky scaffolds with a LEICA VT1200S micro-vibratome with a blade speed of 0.8 mm/s and a vibration amplitude of 1 Hz.

### Water retention measurements

The water retention percentage for each of the PEDOT:PSS scaffold formulations was calculated using the following equation after weighing the scaffolds at dry conditions (directly after freeze dried and annealed) and after immersing the scaffolds in DI water for 24 h. An analytical balance with 0.1 mg resolution was used.

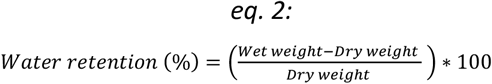

Six different scaffolds were used to calculate the water retention percentage for each formulation. All scaffolds used for the results presented in figure 2 made in the same freeze batch within 98 well-plate. Similar results were obtained from another 2 identically executed experimental runs.

### Young’s modulus

PEDOT:PSS scaffolds were characterized mechanically using Tinius Olsen (Tinius Olsen Ltd) 1-50 kN. With a Tinius Olsen apparatus, the compressive moduli of the dry scaffolds was calculated. The Tinius Olsen had a load cell of 25 N and the compression speed was set at 1 mm min^-1^. Young’s modulus values were calculated from the slope of the linear part of the strain stress curve.

### Scanning electron Microscopy

The microstructure and surface morphology of the scaffolds were analyzed using Scanning Electron Microscopy (Leo Variable pressure SEM, ZEIS GmbH). The samples were mounted on conductive adhesion tape and analyzed at 1 kV power and the beam current was maintained at 50 pA.

### Cell culture

Human adipose-derived stem cells (Lonza - PT-5006) were grown in T-75 flasks in Dulbecco’s modified Eagle medium - high glucose (Sigma – D5671) supplemented with Fetal Bovine Serum (F9665), 2 mM GlutaMax (Gibco-35050061) and 100 U/ml Penicillin-Streptomycin (Gibco-15070063). Prior to cell seeding, scaffolds or devices were immersed into ethanol 70 vol% for at least 1 h for sterilization and then transferred in a sterilized laminar flow hood and washed three times with sterile water and three times with sterile phosphate buffer saline (PBS 1X). The devices were then immersed in cell media and left in a cell incubator at 37 °C/5 % CO_2_ for 2 days to ensure media penetration within the pores. After the 2 days media incubation, a reference EIS spectra was measured and 50000 cells/cm^2^ suspended in 20 ul of complete media were added on top of the devices and incubated for 2 hours to allow the cells to adhere. After the 2 hours period the devices were transferred back in a sterilized laminar flow hood and 280 μL of media were added in the glass wells. The devices were then incubated for several days and fresh media were added every 2 days, after every EIS measurement.

### 3D bioelectronic device fabrication, cell seeding and characterization

ITO substrates were sonicated in acetone and dried with a nitrogen gun. The adhesion layer was deposited on the clean ITO substrates by spin coating a PEDOT:PSS PH 1000 solution with 2 wt% GOPS at 1000 rpm for 45 seconds followed by soft annealing at 90 °C for 30 seconds. Then a wet scaffold slice was gently attached on the PEDOT:PSS adhesion layer and both layers were annealed at 110 °C for 1 h. A micro-patterned kapton tape with a ring opening of 7 mm was attached on the surface of the devices, to accurately define the exposed slice area (0.385 cm^2^). Finally, a glass well was attached on the device using a PDMS as a glue (SYLGARD™ 184 silicone elastomer Kit) and dried overnight at room temperature. After dried, the devices were left in DI water for multiple days and until used. Electrochemical impedance spectroscopy measurements were performed with an Autolab PGSTAT204 by Metrohm AG. An Ag/AgCl/KCL 3M was used as reference and a Pt mesh was used as counter electrode, both immersed in 300 μl of electrolyte (either PBS 1X or cell media). A tungsten pin was used to contact the ITO substrate and served as the working electrode. All electrodes were sterilized with ethanol 70 vol% prior to EIS measurements which were performed in a sterilized laminar flow hood to avoid contamination. The control devices without cells were kept in the same media used to grow hADSCs and kept in an incubator at 37 °C/ 5% CO_2_.

### Live/Dead assay

Cell viability was performed using the LIVE/DEAD_TM_ Viability/Cytotoxicity Kit (Invitrogen) at different time points of the cell culture between day 2 and day 10. This assay permits to assess live and dead cells in vitro by means of two fluorescence molecular probes (i.e. Calcein AM (green, live cells) and Ethidium Homodimer (EthD (red, dead cells), which are representative of intracellular esterase activity and plasma membrane integrity, respectively. The viability assay was performed using 2 μM calcein AM and 4μM EthD-1 in PBS. Scaffold slices were incubated for 45 min at room temperature with the reagents and then mounted on a cover slip for sample observation under the confocal microscope (ZEISS LSM 800). The same approach was followed to address the viability of hADCs within 3D bioelectronic devices.

## Author Contributions

A.S. conceived the research, performed contact angle measurements, fabricated the scaffolds, designed and fabricated the 3D devices, performed water retention measurements, performed EIS measurements, analyzed the results, performed cell culture and wrote the manuscript. J.S.C. performed Young’s modulus measurements, assisted on scaffold fabrication and water retention measurements. J.S.C. and C.M.M. performed SEM measurements. C.B. and C.M.M. performed live/dead assays and confocal microscopy. Z.L. fabricated planar transistors and electrodes and performed OECT and EIS measurements with the assistance of K.K. A.W. assisted on cell culture. C.P. assisted on scaffold fabrication. A.S. and R.M.O. designed the experiments and supervised the work. All authors discussed the results and assisted in manuscript input.

## Acknowledgements

A.S. acknowledges funding from the European Union’s Horizon 2020 research and innovation program under the Marie Skłodowska-Curie grant, MultiStem (No. 895801). J.S. acknowledges funding from the European Union’s Horizon 2020 research and innovation program under the Marie Skłodowska-Curie grant, ICE-METs (No. 842356). K.K. acknowledges funding by the King Abdullah University of Science and Technology (KAUST) Office of Sponsored Research (OSR) under Award No. OSR-2018-CRG7-3709. C.B. and R.M.O. acknowledge funding from the European Research Council (ERC) under the European Union’s Horizon 2020 research and innovation programme (grant agreement no. 723951). The authors also wish to acknowledge funding by the Engineering and Physical Sciences Research Council Centre for Doctoral Training in Sensor Technologies and Applications (EP/L015889/1 to CB).

## Supporting Information

**Figure S1:**
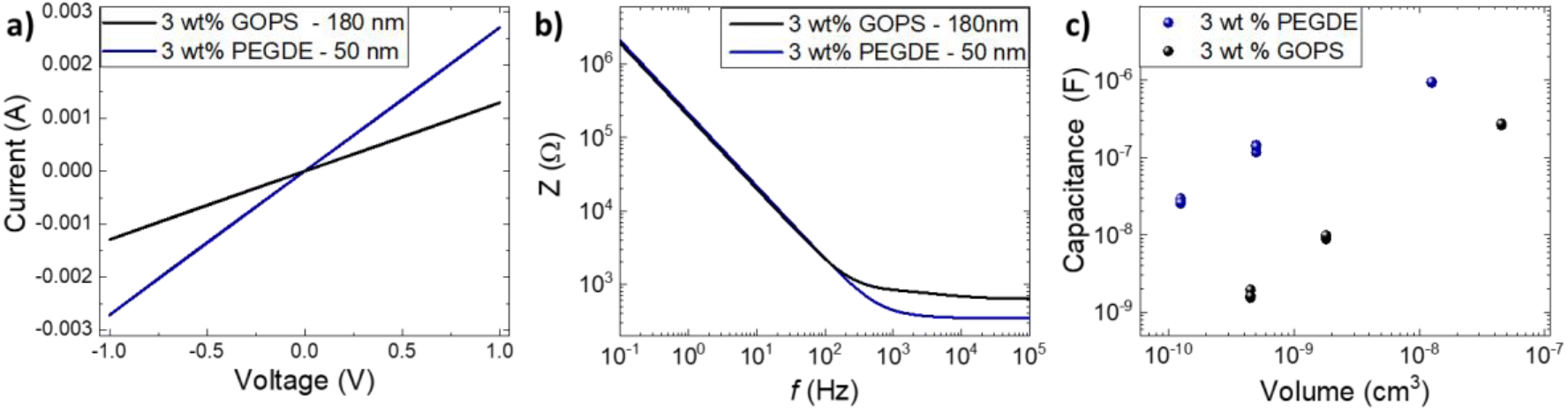
Electrical properties evaluation of PEDOT:PSS films crosslinked with 3 wt% GOPS (black symbols – thickness = 180 nm) and 3 wt % PEGDE (blue symbols thickness = 180 nm). **a)** Current vs voltage characteristics of PEDOT:PSS films casted between two gold electrodes of a fixed distance (100 μm). These data were used to extract the electrical conductivity thin films. **b)** Representative electrochemical impedance spectroscopy measurements of PEDOT:PSS films on micro-fabricated gold square electrodes with side = 500 μm. **c)** the capacitance of PEDOT:PSS films extracted from the impedance at low frequency (i.e 0.1 Hz) for PEDOT:PSS casted on micro-fabricated electrodes with different sizes. All measurements were performed in the aqueous electrolyte PBS 1X.

**Figure S2:**
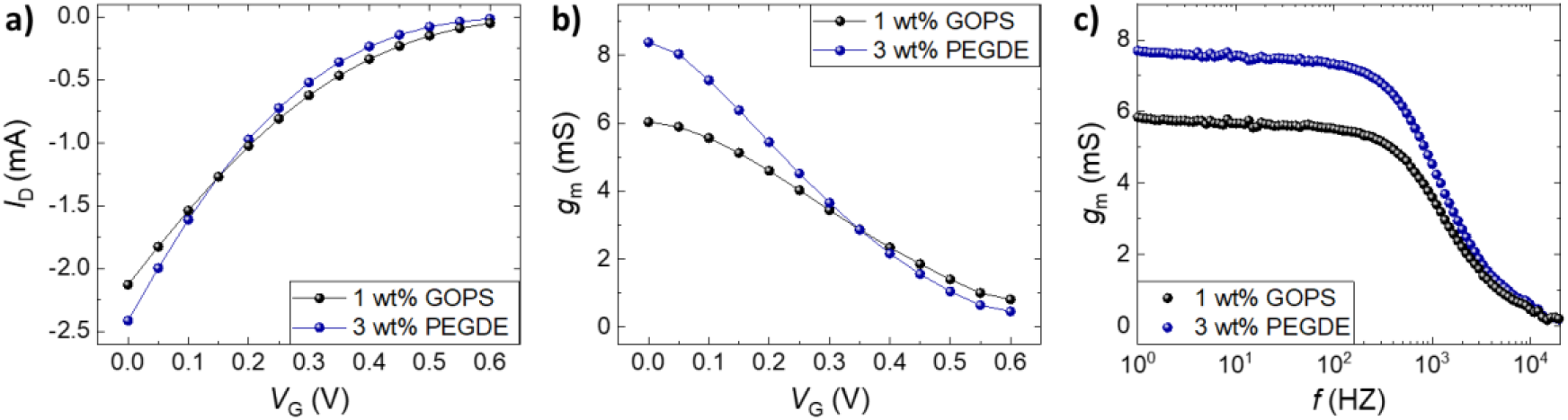
Organic electrochemical transistor characteristics of PEDOT:PSS crosslinked with 3 wt% GOPS (black symbols) and 3 wt % PEGDE (blue symbols). **a)** Transfer curves at *V*_D_ = -0.6V, **b)** transconductance (*g*_m_) at *V*_D_ = -0.6V as a function of gate voltage (*V*_G_) and **c)** transconductance (*g*_m_) at *V*_D_ = -0.6 and *V*_G_ = 0 V as a function of frequency. The cut off frequency is calculated at ∼850 Hz in both cases.

**Figure S3:**
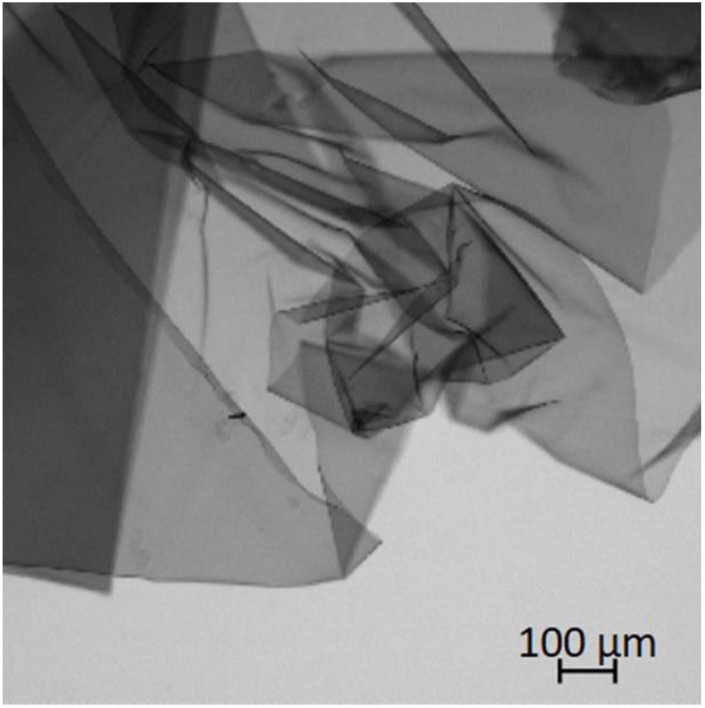
Bright field microscope image of delaminated PEDOT:PSS thin film crosslinked with 3 wt% PEGDE after 5 days immersed in cell media.

**Figure S4:**
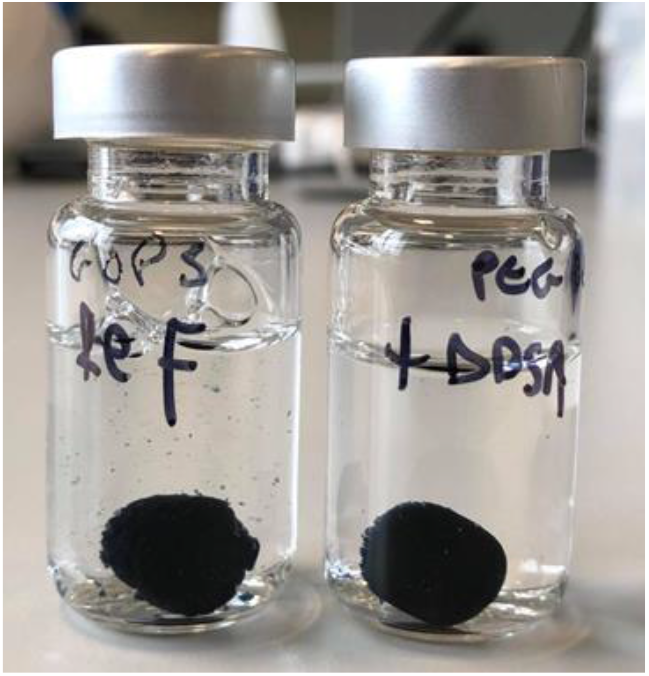
PEDOT:PSS-based scaffolds crosslinked with GOPS 3 wt% (right) and PEGDE 3 wt% (left) immersed in aqueous PBS 1X for more than 6 months.

**Figure S5:**
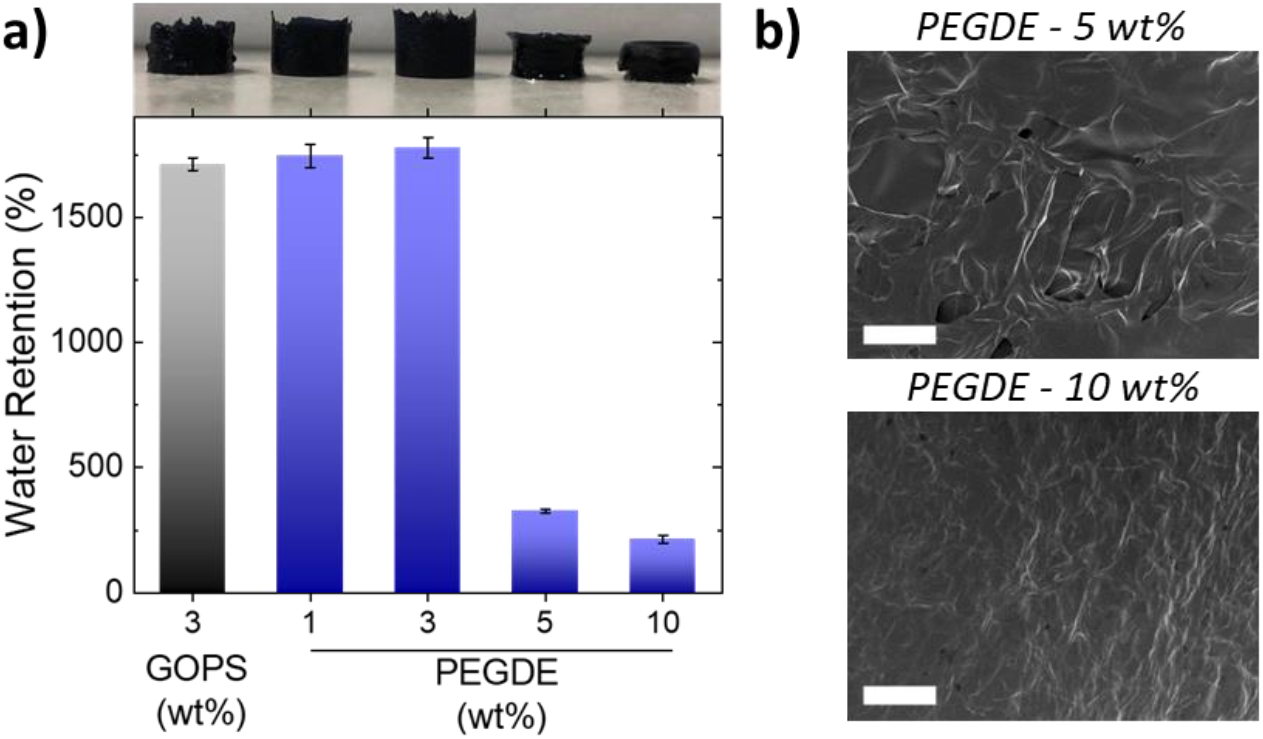
**a)** Water retention ability of PEDOT:PSS-based scaffolds crosslinked with GOPS 3 wt%, PEGDE 1 - 10 wt%. The digital pictures on top of the graph show the scaffolds for the corresponding concentration of crosslinker after been swollen in DI water (N=6). **c)** Scanning electron microscopy measurements of scaffolds prepared with PEGDE 5 wt% and PEGDE 10 wt%. Scale bars = 40 μm.

**Figure S6:**
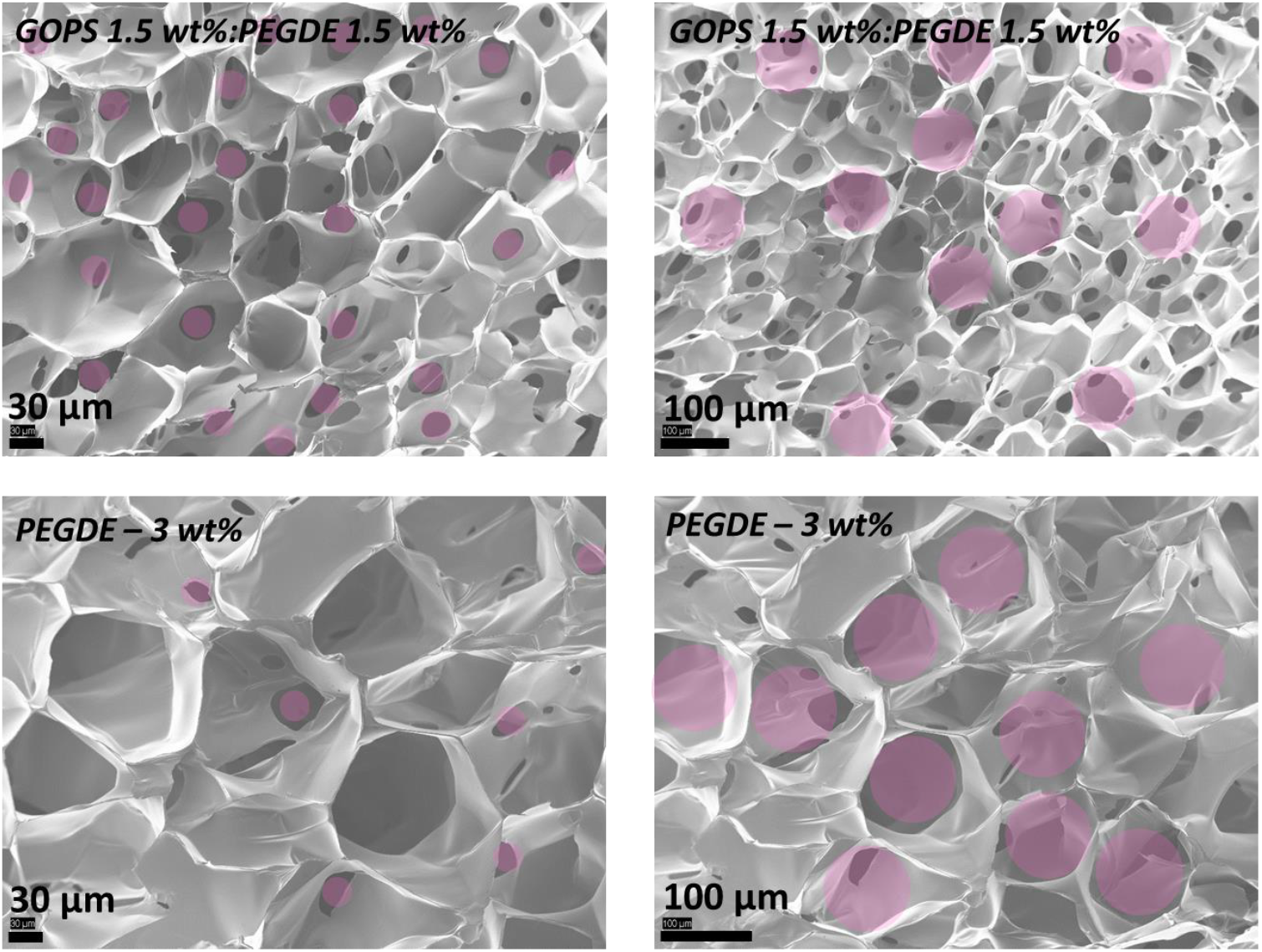
SEM measurements of PEDOT:PSS scaffolds crosslinked with a mixture of GOPS 1.5 wt%:PEGDE 1.5 wt% (top raw) and with PEGDE 3 wt% (bottom raw). The purple circle’s diameter is the same as the scale bar in each image and are used as guides to the eye.

**Figure S7:**
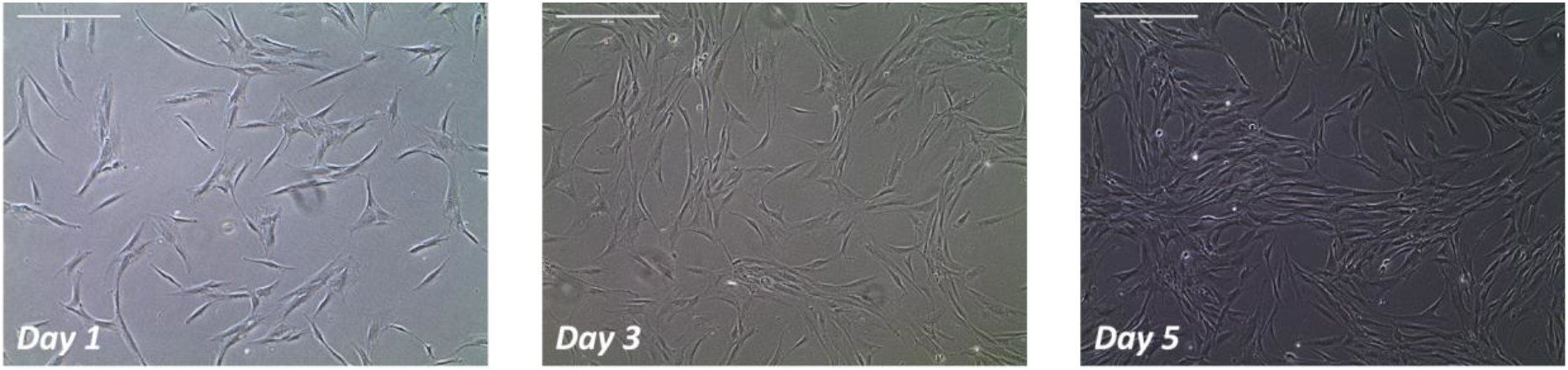
Brightfield images of human adipose derived stem cell cultures in a flat petri dish. Scale bars = 400 μm.

**Figure S8:**
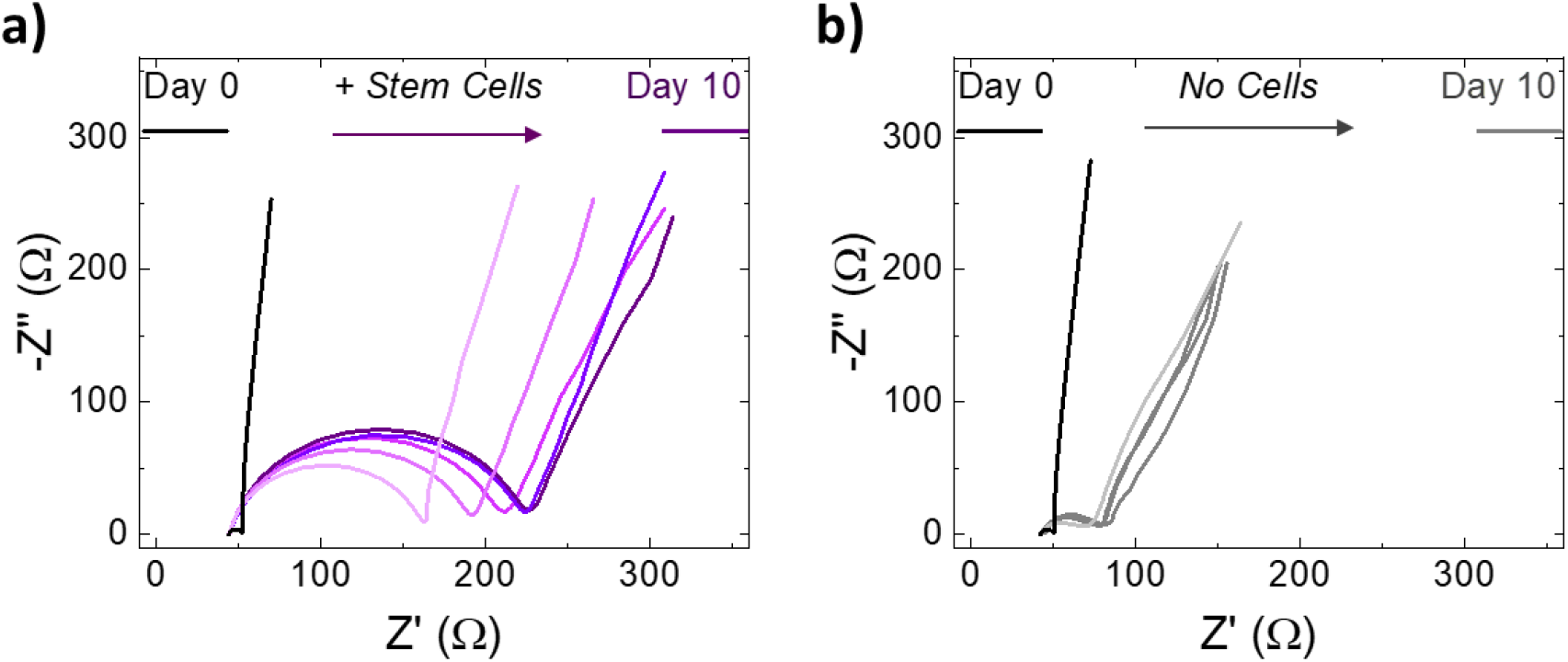
Nyquist plots for **a)** devices seeded with hADSCs and **b)** identically prepared devices not seeded with hADSCs.

## Notes

### Competing Interest Statement

The authors have declared no competing interest.

## References

(1) Zakrzewski, W.; Dobrzyński, M.; Szymonowicz, M.; Rybak, Z. Stem Cells: Past, Present, and Future. Stem Cell Research & Therapy 2019, 10, 68.

(2) Ronaldson-Bouchard, K.; Ma, S. P.; Yeager, K.; Chen, T.; Song, L.; Sirabella, D.; Morikawa, K.; Teles, D.; Yazawa, M.; Vunjak-Novakovic, G. Advanced Maturation of Human Cardiac Tissue Grown from Pluripotent Stem Cells. Nature 2018, 556, 239–243.

(3) Blackiston, D. J.; McLaughlin, K. A.; Levin, M. Bioelectric Controls of Cell Proliferation: Ion Channels, Membrane Voltage and the Cell Cycle. Cell Cycle 2009, 8, 3527–3536.

(4) Henderson, K.; Sligar, A. D.; Le, V. P.; Lee, J.; Baker, A. B. Biomechanical Regulation of Mesenchymal Stem Cells for Cardiovascular Tissue Engineering. Advanced Healthcare Materials 2017, 6, 1700556.

(5) Abagnale, G.; Steger, M.; Nguyen, V. H.; Hersch, N.; Sechi, A.; Joussen, S.; Denecke, B.; Merkel, R.; Hoffmann, B.; Dreser, A.; et al. Surface Topography Enhances Differentiation of Mesenchymal Stem Cells towards Osteogenic and Adipogenic Lineages. Biomaterials 2015, 61, 316–326.

(6) Han, S.-B.; Kim, J.-K.; Lee, G.; Kim, D.-H. Mechanical Properties of Materials for Stem Cell Differentiation. Advanced Biosystems 2020, 4, 2000247.

(7) Pampaloni, F.; Reynaud, E. G.; Stelzer, E. H. K. The Third Dimension Bridges the Gap between Cell Culture and Live Tissue. Nature Reviews Molecular Cell Biology 2007, 8, 839–845.

(8) Nicolas, J.; Magli, S.; Rabbachin, L.; Sampaolesi, S.; Nicotra, F.; Russo, L. 3D Extracellular Matrix Mimics: Fundamental Concepts and Role of Materials Chemistry to Influence Stem Cell Fate. Biomacromolecules 2020, 21, 1968–1994.

(9) Pitsalidis, C.; Ferro, M. P.; Iandolo, D.; Tzounis, L.; Inal, S.; Owens, R. M. Transistor in a Tube: A Route to Three-Dimensional Bioelectronics. Science Advances 2018, 4, 1–10.

(10) Moysidou, C.-M.; Pitsalidis, C.; Al-Sharabi, M.; Withers, A. M.; Zeitler, J. A.; Owens, R. M. 3D Bioelectronic Model of the Human Intestine. Advanced Biology 2021, 5, 2000306.

(11) Inal, S.; Hama, A.; Ferro, M.; Pitsalidis, C.; Oziat, J.; Iandolo, D.; Pappa, A.-M.; Hadida, M.; Huerta, M.; Marchat, D.; et al. Conducting Polymer Scaffolds for Hosting and Monitoring 3D Cell Culture. Advanced Biosystems 2017, 1, 1700052.

(12) Wan, A. M. D.; Inal, S.; Williams, T.; Wang, K.; Leleux, P.; Estevez, L.; Giannelis, E. P.; Fischbach, C.; Malliaras, G. G.; Gourdon, D. 3D Conducting Polymer Platforms for Electrical Control of Protein Conformation and Cellular Functions. Journal of Materials Chemistry B 2015, 3, 5040–5048.

(13) Moysidou, C. M.; Barberio, C.; Owens, R. M. Advances in Engineering Human Tissue Models. Frontiers in Bioengineering and Biotechnology. Frontiers Media S.A. January 2021, p 620962.

(14) Zhang, H. Ice Templating and Freeze-Drying for Porous Materials and Their Applications; John Wiley & Sons, 2018.

(15) Zhang, X.; Li, C.; Luo, Y. Aligned/Unaligned Conducting Polymer Cryogels with Three-Dimensional Macroporous Architectures from Ice-Segregation-Induced Self-Assembly of PEDOT-PSS. Langmuir 2011, 27, 1915–1923.

(16) Jayaram, A. K.; Pitsalidis, C.; Tan, E.; Moysidou, C.-M.; de Volder, M. F. L.; Kim, J.-S.; Owens, R. M. 3D Hybrid Scaffolds Based on PEDOT:PSS/MWCNT Composites. Frontiers in Chemistry 2019, 7.

(17) del Agua, I.; Marina, S.; Pitsalidis, C.; Mantione, D.; Ferro, M.; Iandolo, D.; Sanchez-Sanchez, A.; Malliaras, G. G.; Owens, R. M.; Mecerreyes, D. Conducting Polymer Scaffolds Based on Poly(3,4-Ethylenedioxythiophene) and Xanthan Gum for Live-Cell Monitoring. ACS Omega 2018, 3, 7424–7431.

(18) Håkansson, A.; Han, S.; Wang, S.; Lu, J.; Braun, S.; Fahlman, M.; Berggren, M.; Crispin, X.; Fabiano, S. Effect of (3-Glycidyloxypropyl)Trimethoxysilane (GOPS) on the Electrical Properties of PEDOT:PSS Films. Journal of Polymer Science Part B: Polymer Physics 2017, 55, 814–820.

(19) Dijk, G.; Rutz, A. L.; Malliaras, G. G. Stability of PEDOT:PSS-Coated Gold Electrodes in Cell Culture Conditions. Advanced Materials Technologies 2020, 5, 1900662.

(20) Bidinger, S. L.; Han, S.; Malliaras, G. G.; Hasan, T. Highly Stable PEDOT:PSS Electrochemical Transistors. Applied Physics Letters 2022, 120, 073302.

(21) Solazzo, M.; Krukiewicz, K.; Zhussupbekova, A.; Fleischer, K.; Biggs, M. J.; Monaghan, M. G. PEDOT:PSS Interfaces Stabilised Using a PEGylated Crosslinker Yield Improved Conductivity and Biocompatibility. Journal of Materials Chemistry B 2019, 7, 4811–4820.

(22) del Agua, I.; Mantione, D.; Ismailov, U.; Sanchez-Sanchez, A.; Aramburu, N.; Malliaras, G. G.; Mecerreyes, D.; Ismailova, E. DVS-Crosslinked PEDOT:PSS Free-Standing and Textile Electrodes toward Wearable Health Monitoring. Advanced Materials Technologies 2018, 3, 1700322.

(23) Pitsalidis, C.; Pappa, A.-M.; Boys, A. J.; Fu, Y.; Moysidou, C.-M.; van Niekerk, D.; Saez, J.; Savva, A.; Iandolo, D.; Owens, R. M. Organic Bioelectronics for In Vitro Systems. Chemical Reviews 2021.

(24) ElMahmoudy, M.; Inal, S.; Charrier, A.; Uguz, I.; Malliaras, G. G.; Sanaur, S. Tailoring the Electrochemical and Mechanical Properties of PEDOT:PSS Films for Bioelectronics. Macromolecular Materials and Engineering 2017, 302, 1600497.

(25) del Agua, I.; Mantione, D.; Ismailov, U.; Sanchez-Sanchez, A.; Aramburu, N.; Malliaras, G. G.; Mecerreyes, D.; Ismailova, E. DVS-Crosslinked PEDOT:PSS Free-Standing and Textile Electrodes toward Wearable Health Monitoring. Advanced Materials Technologies 2018, 3, 1700322.

(26) Iandolo, D.; Sheard, J.; Karavitas Levy, G.; Pitsalidis, C.; Tan, E.; Dennis, A.; Kim, J. S.; Markaki, A. E.; Widera, D.; Owens, R. M. Biomimetic and Electroactive 3D Scaffolds for Human Neural Crest-Derived Stem Cell Expansion and Osteogenic Differentiation. MRS Communications 2020, 10, 179–187.

(27) Guex, A. G.; Puetzer, J. L.; Armgarth, A.; Littmann, E.; Stavrinidou, E.; Giannelis, E. P.; Malliaras, G. G.; Stevens, M. M. Highly Porous Scaffolds of PEDOT:PSS for Bone Tissue Engineering. Acta Biomaterialia 2017, 62, 91–101.

(28) Zhou, X.; Holsbeeks, I.; Impens, S.; Sonnaert, M.; Bloemen, V.; Luyten, F.; Schrooten, J. Noninvasive Real-Time Monitoring by AlamarBlue(®) during in Vitro Culture of Three-Dimensional Tissue-Engineered Bone Constructs. Tissue Eng Part C Methods 2013, 19, 720–729.

(29) Park, H. E.; Kim, D.; Koh, H. S.; Cho, S.; Sung, J.-S.; Kim, J. Y. Real-Time Monitoring of Neural Differentiation of Human Mesenchymal Stem Cells by Electric Cell-Substrate Impedance Sensing. Journal of Biomedicine and Biotechnology 2011, 2011, 485173.

(30) Ryan, A. J.; Kearney, C. J.; Shen, N.; Khan, U.; Kelly, A. G.; Probst, C.; Brauchle, E.; Biccai, S.; Garciarena, C. D.; Vega-Mayoral, V.; et al. Electroconductive Biohybrid Collagen/Pristine Graphene Composite Biomaterials with Enhanced Biological Activity. Advanced Materials 2018, 30, 1706442.

(31) Savva, A.; Hama, A.; Herrera-López, G.; Gasparini, N.; Migliaccio, L.; Kawan, M.; Steiner, N.; McCulloch, I.; Baran, D.; Fiumelli, H.; et al. Photo-Electrochemical Stimulation of Neurons with Organic Donor-Acceptor Heterojunctions. bioRxiv 2022, 2022.02.17.480608.

(32) Rivnay, J.; Leleux, P.; Ferro, M.; Sessolo, M.; Williamson, A.; Koutsouras, D. A.; Khodagholy, D.; Ramuz, M.; Strakosas, X.; Owens, R. M.; et al. High-Performance Transistors for Bioelectronics through Tuning of Channel Thickness. Science Advances 2015, 1, e1400251.

(33) Barberini, D. J.; Freitas, N. P. P.; Magnoni, M. S.; Maia, L.; Listoni, A. J.; Heckler, M. C.; Sudano, M. J.; Golim, M. A.; da Cruz Landim-Alvarenga, F.; Amorim, R. M. Equine Mesenchymal Stem Cells from Bone Marrow, Adipose Tissue and Umbilical Cord: Immunophenotypic Characterization and Differentiation Potential. Stem Cell Research & Therapy 2014, 5, 25.

(34) Bianchi, M.; Carli, S.; di Lauro, M.; Prato, M.; Murgia, M.; Fadiga, L.; Biscarini, F. Scaling of Capacitance of PEDOT:PSS: Volume vs. Area. Journal of Materials Chemistry C 2020, 8, 11252–11262.

(35) Proctor, C. M.; Rivnay, J.; Malliaras, G. G. Understanding Volumetric Capacitance in Conducting Polymers. Journal of Polymer Science Part B: Polymer Physics 2016, 54, 1433–1436.

(36) Volkov, A. v; Wijeratne, K.; Mitraka, E.; Ail, U.; Zhao, D.; Tybrandt, K.; Andreasen, J. W.; Berggren, M.; Crispin, X.; Zozoulenko, I. v. Understanding the Capacitance of PEDOT:PSS. Advanced Functional Materials 2017, 27, 1700329.

(37) Duc, C.; Vlandas, A.; Malliaras, G. G.; Senez, V. Wettability of PEDOT:PSS Films. Soft Matter 2016, 12, 5146–5153.

